# Potential antigenic mismatch of the H3N2 component of the 2019 Southern Hemisphere influenza vaccine

**DOI:** 10.1101/681734

**Authors:** Sigrid Gouma, Madison Weirick, Scott E. Hensley

**Affiliations:** Department of Microbiology, Perelman School of Medicine, University of Pennsylvania, Philadelphia, PA, USA

**Keywords:** Influenza virus, hemagglutinin, influenza vaccine, H3N2

## Abstract

Here, we find that the egg-adapted H3N2 component of the 2019 Southern Hemisphere influenza vaccine elicits an antibody response in ferrets that is highly focused on antigenic site A of hemagglutinin. This is potentially problematic since most H3N2 viruses currently circulating in the Southern Hemisphere possess antigenic site A substitutions.

---

Human egg-adapted 3c2.A H3N2 influenza vaccine strains possess adaptive substitutions that result in the loss of a key glycosylation site within antigenic site B of hemagglutinin (HA) [1]. We previously demonstrated that the 2016-2017 Northern Hemisphere egg-adapted H3N2 vaccine strain (A/Hong Kong/4801/2014) elicits HA antigenic site B antibodies that bind to the vaccine strain but not to circulating 3c2.A H3N2 strains [1]. This antigenic mismatch has been associated with low vaccine effectiveness (VE) during recent H3N2-dominated influenza seasons [2, 3].

3c2.A viruses recently diversified into several subgroups (**Figure 1A**), and a new H3N2 vaccine strain, A/Switzerland/8060/2017 (3c2.A2 subgroup), was recommended for the 2019 Southern Hemisphere vaccine formulation [4]. Similar to previous egg-adapted 3c2.A H3N2 vaccine strains, the HA of A/Switzerland/8060/2017 possesses a T160K HA substitution that results in the loss of a glycosylation site in antigenic site B of HA (**Figure 1B**). Interestingly, data from the World Health Organization indicate that antibodies elicited by the egg-adapted A/Switzerland/8060/2017 strain are less focused towards HA antigenic site B compared to antibodies elicited by previous 3c2.A H3N2 egg-adapted vaccine strains [4]. To determine if the A/Switzerland/8060/2017 strain is antigenically matched to H3N2 strains currently circulating in the Southern Hemisphere, we completed antigenic analyses using sera isolated from ferrets exposed to either wild-type or egg-adapted A/Switzerland/8060/2017 strains.

**Figure 1.**
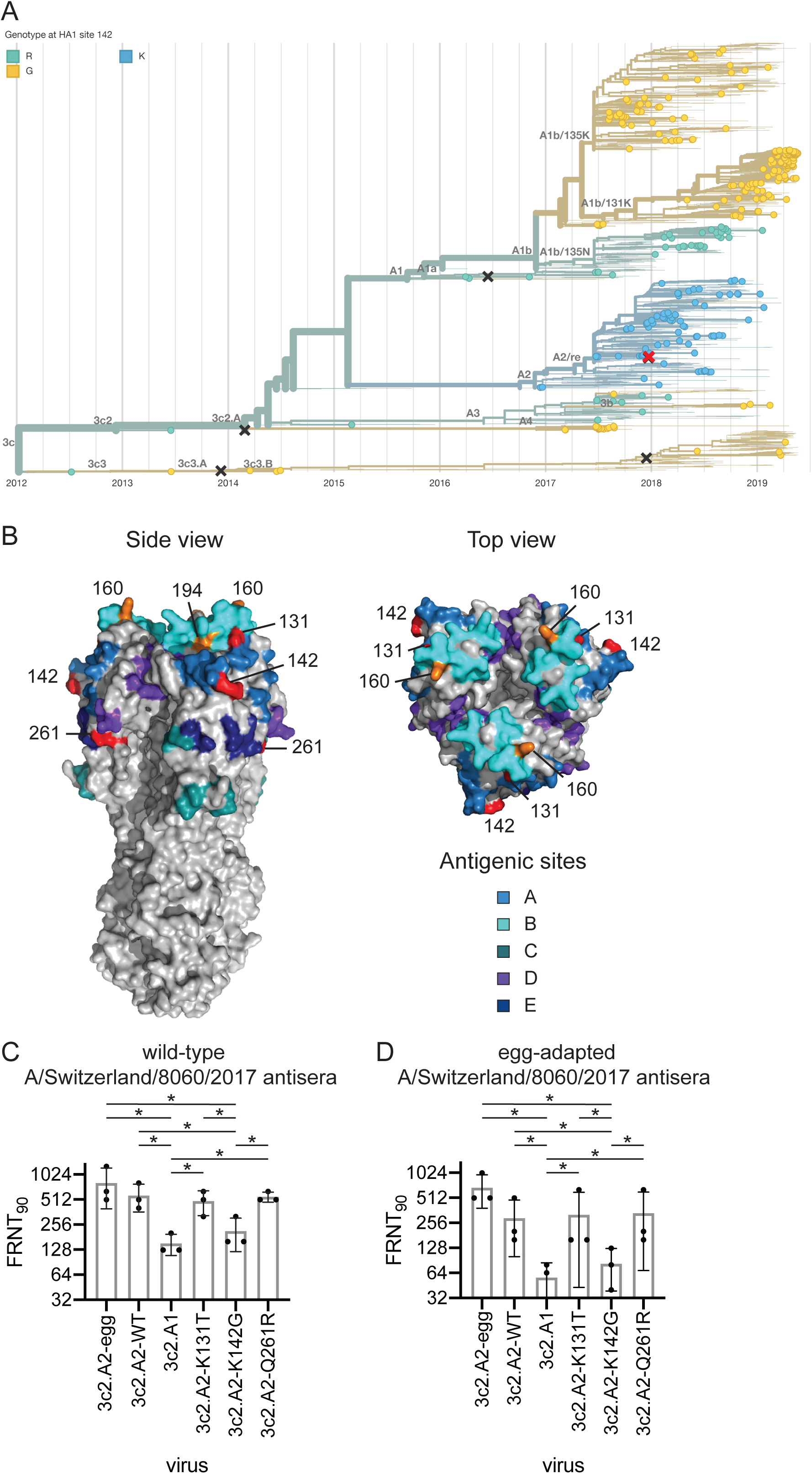
Potential antigenic mismatch of the H3N2 component of the 2019 Southern Hemisphere influenza vaccine. (A) Phylogenetic tree showing the diversification of 3c2.A viruses and the circulation of 3c2.A1 viruses in Oceania (n=306) and South America (n=227). The phylogenetic tree was retrieved from www.nextrain.org on June 21, 2019 [6]. H3N2 vaccine components used since 2015 are indicated with ‘x’ marks. The red ‘x’ mark indicates the H3N2 component of the 2019 Southern Hemisphere influenza vaccine A/Switzerland/8060/2017 (clade 3c2.A2). Sequences are colored by their residue at site 142 in the HA protein. (B) Crystal structure of the HA trimer of A/Victoria/361/2011 (PDB accession code 4O5I) with the antigenic sites indicated as previously described [10, 11]. Egg-adapted substitutions at positions 160 and 194 are shown in orange. We introduced substitutions in HA residues indicated in red for antigenic testing. (C-D) FRNTs were performed using sera from ferrets recovering from infection with wild-type A/Switzerland/8060/2017 or egg-adapted A/Switzerland/8060/2017 antisera. * p<0.05.

## Methods

### Infection of ferrets with influenza virus

We used reverse-genetics [5] to create viruses with the wild-type or egg-adapted A/Switzerland/8060/2017 HA. The egg-adapted HA possessed T160K and L194P substitutions in HA antigenic site B relative to the wild-type HA (**Figure 1B**). We rescued viruses using A/Puerto Rico/8/1934 internal genes to allow efficient viral growth in culture. Ferrets were infected with 3,000 foci forming units (FFU) of virus with either HA, and then bled 35 days later. All animal experiments were completed under an Institutional Animal Care and Use Committee-approved protocol at Nobel Life Sciences.

### Foci Reduction Neutralization Tests (FRNTs)

Serum samples were treated with receptor-destroying enzyme (Denka Seiken) followed by heat-inactivation at 55°C for 30 minutes prior to testing in FRNTs. FRNTs were performed as previously described [1] using ferret sera and 3c2.A2 viruses with wild-type and egg-adapted HAs. We also completed assays with a 3c2.A1 virus (wild-type A/Singapore/INFIMH-16-0019/2016) and 3c2.A2 viruses with single HA substitutions. Each sample was tested in three independent experiments and geometric mean titers of the replicates were used for analysis.

### Statistical methods

Log_2_-transformed antibody titers were compared using RM one-way ANOVA corrected for multiple comparisons (Bonferroni method). Statistical analysis was performed using Prism version 7.

## Results

Ferrets infected with the wild-type A/Switzerland/8060/2017 strain mounted antibody responses that neutralized wild-type and egg-adapted 3c2.A2 viruses equivalently (**Figure 1C**). Ferrets infected with the egg-adapted A/Switzerland/8060/2017 strain mounted antibody responses that efficiently neutralized egg-adapted 3c2.A2 virus and moderately neutralized wild-type 3c2.A2 virus (**Figure 1D**). Importantly, antibodies elicited by the wild-type and egg-adapted A/Switzerland/8060/2017 strains poorly neutralized the 3c2.A1 virus (**Figure 1D**). This is important since 3c2.A1 viruses are currently circulating at high levels in parts of the Southern Hemisphere [6]. These data suggest that antibodies elicited by the A/Switzerland/8060/2017 strain are minimally affected by antigenic site B HA substitutions that distinguish wild-type and egg-adapted 3c2.A2 strains. This indicates that most antibodies elicited in ferrets by wild-type and egg-adapted A/Switzerland/8060/2017 strains target other HA epitopes that are mismatched between 3c2.A2 and 3c2.A1 viruses.

To further define the specificity of antibodies elicited by wild-type and egg-adapted A/Switzerland/8060/2017 strains, we completed additional FRNTs with a panel of 3c2.A2 viruses with different HA substitutions. We tested 3c2.A2 viruses that possess each of the HA substitutions (K131T, K142G, and Q261R) that distinguish 3c2.A2 viruses from other 3c2.A clades. Residue 142 is located in antigenic site A, residue 131 is located at the interface of antigenic sites A and B, and residue 261 is located in close proximity to antigenic site E (**Figure B**). Antibodies elicited by both the wild-type and egg-adapted A/Switzerland/8060/2017 strains efficiently neutralized 3c2.A2 viruses with either the K131T or Q261R HA substitutions but had reduced reactivity to the 3c2.A2 virus with the K142G HA substitution (**Figures 1C-D**). These data indicate that unlike past 3c2.A egg-adapted strains that elicit a heavily focused antigenic site B HA response [1], the egg-adapted A/Switzerland/8060/2017 strain elicits antibody responses that are more directed towards epitopes involving residue 142 that is located in antigenic site A of HA.

## Discussion

It is interesting that the egg-adapted A/Hong Kong/4801/2014 strain elicits an antibody response focused towards HA antigenic site B [1], whereas we find that the egg-adapted A/Switzerland/8060/2017 strain elicits an antibody response focused towards HA antigenic site A. We previously demonstrated that HA antigenic site A of a 1995 H3N2 strain was immunodominant, only if the HA possessed an N145K substitution within antigenic site A [7]. Similarly, Santos *et al.* found that antigenic site A became immunodominant when K145 was introduced into a panel of distantly related H3 HAs [8]. The immune basis for these apparent immunodominance shifts are unknown, but it is possible that the presence of Lys in HA antigenic site A increases immunogenicity. Consistent with this idea, the egg-adapted A/Switzerland/8060/2017 HA possesses K142, whereas the egg-adapted A/Hong Kong/4801/2014 HA possesses R142 and 3c2.A1 HAs possess G142.

There are several limitations of our study. Our study evaluated antibody responses elicited in ferrets, not humans. Studies from our laboratory and others [9] clearly demonstrate that the specificity of influenza virus antibodies elicited in ferrets and humans can differ because most humans have complex immune histories, whereas ferrets do not. It is also worth noting that we tested serum from ferrets that were intranasally infected with virus rather than vaccinated. Ferret serum used for antigenic analyses are typically prepared in this manner [4] but it is possible that vaccinations with the egg-adapted A/Switzerland/8060/2017 strain elicit antibodies of different specificities compared to infections.

The implications of these experiments are obvious: the egg-adapted H3N2 component of the 2019 Southern Hemisphere vaccine might elicit suboptimal antibody responses that react poorly to HA site A of 3c2.A1 viruses, which are currently dominating in parts of the Southern Hemisphere. Vaccination is obviously still recommended because it is unclear if 3c2.A1 viruses will continue to dominate circulation and it is unknown if vaccine effectiveness will be impacted by the potential antigenic mismatch identified in our study. Further, the H1N1 and influenza B virus components of the 2019 Southern Hemisphere vaccine are likely well-matched to circulating strains and thus likely to provide good protection.

## References

1. Zost SJ, Parkhouse K, Gumina ME, et al. Contemporary H3N2 influenza viruses have a glycosylation site that alters binding of antibodies elicited by egg-adapted vaccine strains. Proc Natl Acad Sci 2017; 114:12578–12583.

2. Zimmerman RK, Nowalk MP, Chung J, et al. 2014–2015 influenza vaccine effectiveness in the United States by vaccine type. Clin Infect Dis 2016; 63:1564–1573.

3. Rolfes MA, Flannery B, Chung J, et al. Effects of influenza vaccination in the United States during the 2017–2018 influenza season. Clin Infect Dis 2019; DOI:10.1093/cid/ciz075.

4. World Health Organization. Recommended composition of influenza virus vaccines for use in the 2019 southern hemisphere influenza season.

5. Chambers BS, Parkhouse K, Ross TM, Alby K, Hensley SE. Identification of hemagglutinin residues responsible for H3N2 antigenic drift during the 2014–2015 influenza season. Cell Rep 2015; 12:1–6.

6. Hadfield J, Megill C, Bell SM, et al. Nextstrain: real-time tracking of pathogen evolution. Bioinformatics 2018; 34:4121–4123.

7. Li Y, Bostick DL, Sullivan CB, et al. Single hemagglutinin mutations that alter both antigenicity and receptor binding avidity influence influenza virus antigenic clustering. J Virol 2013; 87:9904–9910.

8. Santos JJS, Abente EJ, Obadan AO, et al. Plasticity of amino acid residue 145 near the receptor binding site of H3 swine influenza A viruses and its impact on receptor binding and antibody recognition. J Virol 2019; 93:e01413–18.

9. Cobey S, Hensley SE. Immune history and influenza virus susceptibility. Curr Opin Virol 2017; 22:105–111.

10. Wiley DC, Skehel JJ. The Structure and function of the hemagglutinin membrane glycoprotein of influenza virus. Annu Rev Biochem 1987; 56:365–394.

11. Popova L, Smith K, West AH, et al. Immunodominance of antigenic site B over site A of hemagglutinin of recent H3N2 influenza viruses. PLoS One 2012; 7:e41895.

